# Subgoal- and Goal-Related Prediction Errors in Medial Prefrontal Cortex

**DOI:** 10.1101/245829

**Authors:** José J. F. Ribas Fernandes, Danesh Shahnazian, Clay B. Holroyd, Matthew M. Botvinick

## Abstract

A longstanding view of the organization of human and animal behavior holds that behavior is hierarchically organized, meaning that it can be understood as directed towards achieving superordinate goals through subordinate goals, or subgoals. For example, the superordinate goal of making coffee can be broken down as accomplishing a series of subgoals, namely boiling water, grinding coffee, pouring cream, etc.

Learning and behavioral adaptation depend on prediction-error signals, which have been observed in ventral striatum (VS) and medial prefrontal cortex (mPFC). In past work, we have shown that prediction error signals (PEs) can be linked not only to superordinate goals, but also to subgoals.

Here we present two functional magnetic resonance imagining experiments that replicate and extend these findings. In the first experiment, we replicated the finding that mPFC signals subgoal-related PEs, independently of goal PEs. Together with our past work, this experiment reveals that BOLD responses to PEs in mPFC are unsigned. In the second experiment, we showed that when a task involves both goal and subgoal PEs, mPFC shows only goal-related PEs, suggesting that context or attention can strongly impact hierarchical PE coding. Furthermore, we observed a dissociation between the coding of PEs in mPFC and VS. These experiments suggest that the mPFC selectively attends to information at different levels of hierarchy depending on the task context.

## Introduction

Learning and behavioral adaptation depend on prediction-error signals (PE)— signals that are generated when the agent’s expectation about future events are violated. The neural correlates of signed PEs have been observed across a variety of experimental paradigms in Ventral Striatum (VS) and medial Prefrontal Cortex (mPFC) (Dolan & Dayan, 2013; Hyman, Holroyd, & Seamans, 2017; Niv, 2009; Roesch, Esber, Li, Daw, & Schoenbaum, 2012). Most of these studies have examined tasks involving one step choice. However, human and animal decision making often involves a sequence of steps (Botvinick, Niv, & Barto, 2009; Lashley, 1951). In multi-step behavior, goals can be parsed into sub-goals, and thus PEs can exist to both goals and sub-goals (Botvinick et al., 2009).

In past work, we have shown that PE signals in both VS and mPFC reflect not only task goals, but also task sub-goals (VS: Diuk, Tsai, Wallis, Botvinick, & Niv, 2013; mPFC: Ribas-Fernandes et al., 2011). Ribas-Fernandes et al. (2011) used a multi-step navigation task, with explicit subgoal and goal states, in which distance to subgoal and distance to goal could be manipulated independently. Unexpectedly, the authors increased the distance to the subgoal, which made its attainment more difficult, without affecting the attainment of goal. This way, the authors elicited negative subgoal-related PEs without goal-related prediction errors. The authors observed an increase in BOLD signal in mPFC, and anterior insula, using functional magnetic resonance imaging (fMRI). This was matched with a frontocentral negativity observed in a parallel electroencephalogram (EEG) experiment, consistent with an involvement of mPFC (Miltner et al., 1997; Miltner, Braun, & Coles, 1997). Further analysis showed that the effects could not be attributed to perceptual and motor elements of the task. Diuk et al. (2013) further explored the correlates of both subgoal and goal-related PEs, using a casino-like three step choice task. In this study, only VS was shown to respond to subgoal and goal-related PEs and the authors failed to observe an effect in mPFC.

Moreover, if we take results in Ribas Fernandes et al. at its face value, three questions are left unsettled: Which brain areas encode positive subgoal-related PEs? Which brain areas encode goal-related PEs elicited using the navigation task? How do the correlates of goal-related PEs with the navigation task compare with PEs elicited with standard methods (e.g., monetary deviations)? Here, using the navigation paradigm of Ribas-Fernandes et al., we examine these three questions in two functional magnetic resonance imagining experiments, eliciting positive subgoal-related PEs (Experiment I), and positive and negative, goal and subgoal-related PEs (Experiment II). Answering these questions will allow us to more directly compare the findings of Ribas-Fernandes et al. (2011) and Diuk et al. (2013). We use a combination of whole brain analysis and region of interest (ROI) analysis to investigate the neural correlates of subgoal-related PE and goal-related PE.

## Experiment I: An fMRI Examination of Positive Subgoal-Related PEs

### Materials and Methods

**Participants**. Participants were recruited from the Princeton University community and all gave their informed consent. 30 participants were recruited (ages 18–25, *M* = 20.5 years, 11 males, all were right-handed). All participants received monetary compensation at a departmental standard rate. To further encourage performance, participants also received a small monetary bonus based on task performance.

**Task and procedure. *Task rationale***. We used a hierarchical multi-step spatial paradigm (see Figure 1A; Ribas-Fernandes, et al., 2011). On each trial, human participants had to pick up an envelope and deliver it to a house, using a joystick to guide a truck. Each joystick movement displaced the truck by a fixed distance. We assume that participants represent the task hierarchically: meaning that they construe delivery to the house as the top-level goal, or what we refer as task level, and acquisition of the envelope as a subgoal, or subtask level.

**Figure 1.**
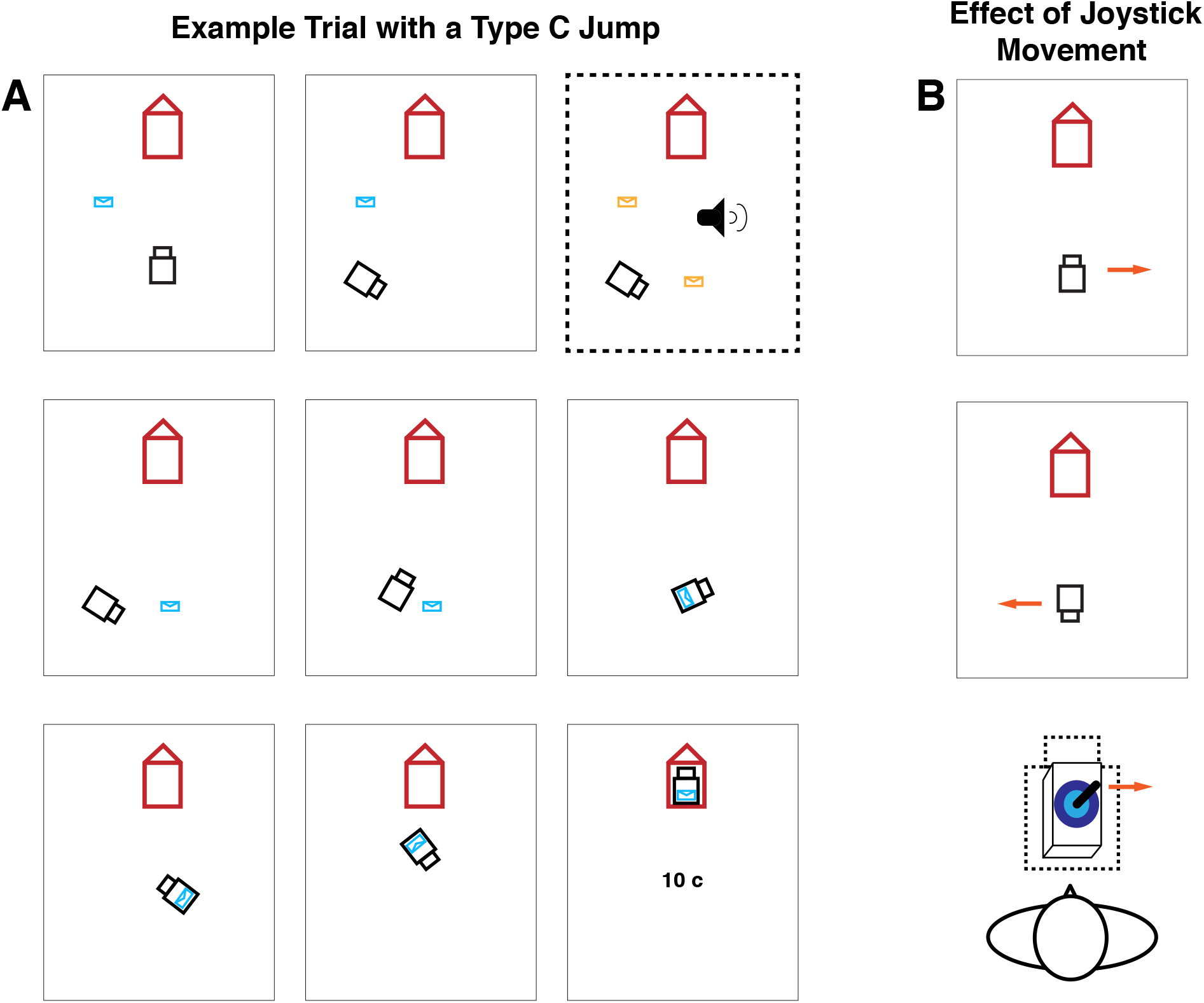
Hierarchical delivery task. *A*. In this task participants had to move a truck, using a joystick, to pick up an envelope and deliver it to a house. Each joystick movement displaced the truck by 50 pixels (note that distance between start point and envelope was 395 pixels). However, after each movement, the orientation of the truck would change randomly. In two thirds of the trials, the envelope would jump to a new location before the truck had reached it, signaled by a beep and a forced pause for 900 ms (see panel bordered by the dashed line). In the remaining third of trials only the beep and the pause would happen. After delivering the envelope to the house participants would be rewarded with 10c. *B*. In order to ensure that each step would be cognitively effortful, the effect of joystick movements was contingent on the orientation of the truck relative to the screen.For example, if the truck were facing downwards, as illustrated in the bottom panel, a rightward movement would displace the truck to the left of the screen.

Importantly, after each movement the truck would randomly change its orientation (Figure 1A). To move the truck in a desired direction, the angle of the joystick had to compensate for the random deviation of the truck (Figure 1B). Given that each movement required sensorimotor coordination, we expected that participants preferred traversing shorter distances in delivering the envelope. Indeed, in an independent behavioral assay, where participants could choose between two envelope delivery trajectories, differing in overall distance to the goal, they would overwhelmingly prefer the shorter route (Ribas-Fernandes et al., 2011).

Because of the spatial nature of the paradigm, it is possible to change the distance to the subgoal (start–envelope), without changing the overall distance (start–envelope– house) (Figure 2). Geometrically, all points on an ellipse with foci on the truck and the house have the same overall distance from start to envelope to house, though different distances to the envelope.

**Figure 2.**
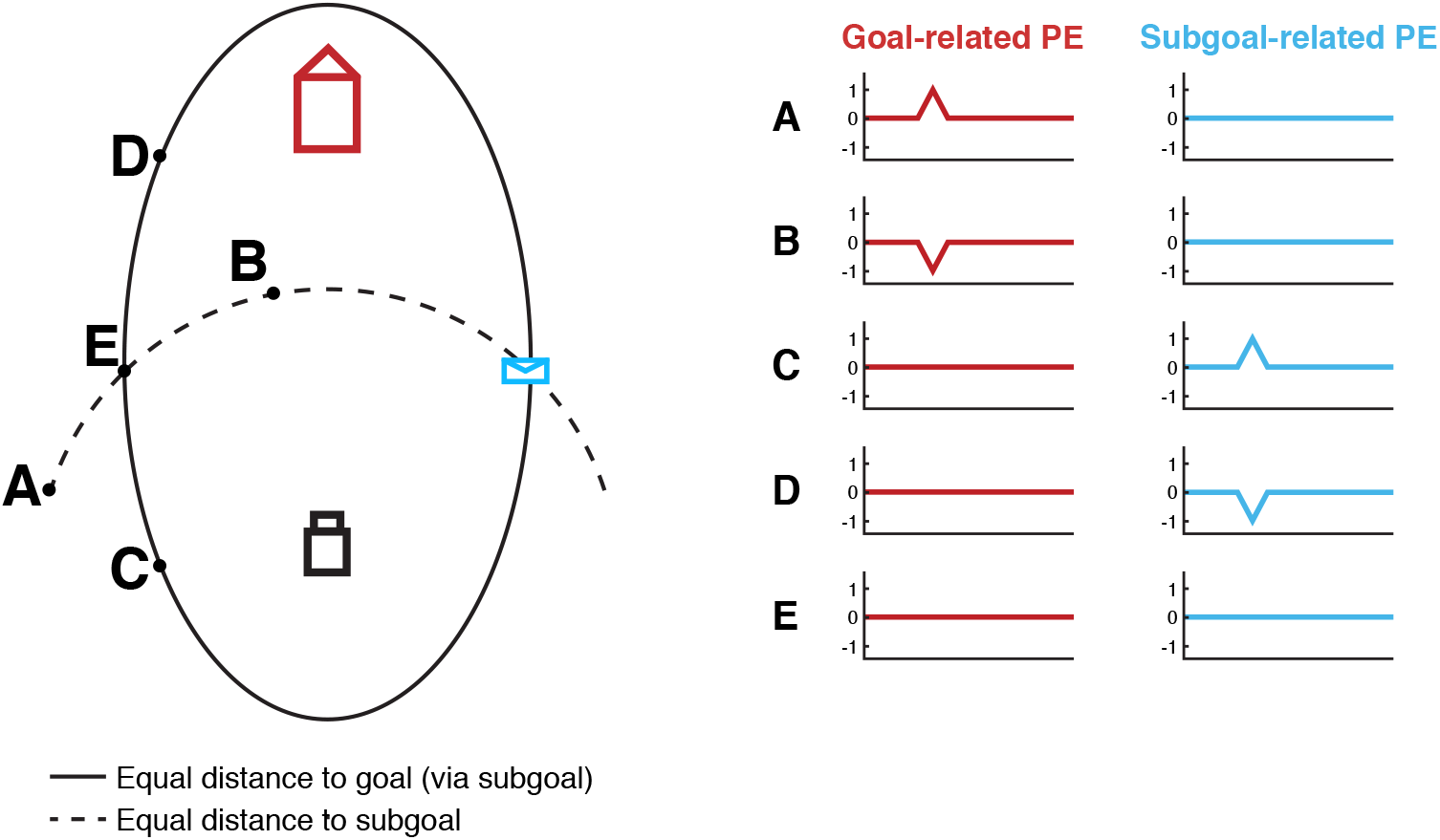
Different types of PEs induced by jumps of the envelope to different screen locations. Left view is task display and underlying geometry of the delivery task. Jumps to points on the solid black line, including D, E, and C, preserve the overall distance to the goal (start-to-envelope summed with envelope-to-house). Therefore, points on the solid black line only differ in their distance to the subgoal. Right view shows PE signals generated in each category of jump event. In Experiment I, the envelope would jump to locations C on a third of the trials (triggering a positive subgoal-related PE), to location E in a third of the trials, and remain in the same place for a third of the trials. In Experiment II, the envelope would jump to locations that would trigger both a goal-related PE and a subgoal-related PE (not shown) on two-thirds of the trials, and remain in the same place in a third of the trials.

We hypothesize that unexpected changes to the distance to the goal elicit goal-related prediction errors. An unexpected increase in distance to the house should elicit a negative goal-related prediction error, whereas a decrease in distance elicits a positive goal-related prediction error. The same follows for displacements of the envelope, namely increases in the distance to the subgoal elicit a negative subgoal-related prediction error, and a decrease in distance to the subgoal/envelope elicits a positive subgoal-related prediction error. In this experiment, we elicited positive subgoal-related prediction errors (see jump C in Figure 2). Additionally, to control for perceptual and motor changes associated with an envelope jump, we also introduced jumps that preserved the distance to both subgoal and goal (see jump to location E).

***Procedure***. The computerized task was coded using MATLAB (The MathWorks, Natick, MA, USA) and the MATLAB Psychophysics toolbox, version 3 (Brainard, 1997). An MR-compatible joystick was used for the scanning part of the task (MagConcept, Redwood City, CA, USA), whereas a regular joystick was used for trials outside the scanner (Logitech International, Romanel-sur-Marges, Switzerland).

On each trial, the starting positions of the icons (truck, envelope, house) were vertices in a triangle with fixed distances and angles. The actual positions were random rotations or reflections of the following triangle: truck, 0, 200, envelope, 151, −165, and house, 0, −200 (x,y coordinates in pixels, referenced to the center of a 1024 × 768 pixels screen). Therefore, the distance between the start point and the envelope was 395 pixels, and the distance between the envelope and the house was 365 pixels, totaling 760 pixels. We should stress that, as mentioned before, actual positions could vary due to the random rotations and reflections, but the distances and angles between icons were preserved.

Each joystick movement displaced the truck by 50 pixels. The direction of the displacement was a function of the truck’s angle with the screen’s vertical axis and the angle of the hand movement, inputted through the joystick, relative to center front of the joystick (Figure 1B). After each displacement, the angle between the truck and the screen’s vertical axis, was changed randomly. Therefore, participants had to adjust the angle of their hand movements on each step, to move the truck in the intended direction.

On every trial, after the first, second or third joystick movement, a brief tone occurred and the envelope flashed for 900 ms, during which joystick movements were ignored (Figure 1A), we hereafter refer to this event as a pause event. On one third of the pause events, the envelope remained in its original location (no-jump condition). On the remaining trials, at the onset of the tone, the envelope jumped to a new location (jump condition). We will use the term pause event for the combination of the no-jump and jump condition. In half of the trials in the jump condition, the distance between the envelope’s new position and the truck position was unchanged by the jump (case E in Figure 2). On the remaining third, a type C jump would happen, the destination of the envelope was chosen such that the distance between truck and envelope always decreased to 120 pixels while the overall path length to the goal (house) was left unchanged. Participants were told that the envelope sometimes stayed in the same place, and sometimes it jumped, with no mention of the distinction between jumps to E *vs*. C.

After the jump, participants were required to navigate towards the new location of the subgoal. When the truck passed within 30 pixels of the envelope, the envelope moved to the truck and remained there for the subsequent moves (Figure 1A). When the truck with the envelope passed within 30 pixels of the house, the image of the truck and the envelope appeared in the house. This image was displayed for 200 ms.

At the completion of each trial (which required on average 17.16 steps or joystick movements), subjects were rewarded with 10 cents (US dollars). This was indicated by a screen displaying “10 c” for 500 ms. Immediately following this, a fixation cross appeared for 2500 ms, followed by onset of the next trial, signaled by the appearance of a new spatial arrangement of icons.

Given that the task requires complex sensory-motor coordination, participants practiced the task prior to functional data acquisition. Practice consisted of fifteen minutes outside the scanner, followed by an eight-minute session inside the scanner during structural scan acquisition.

Inside the scanner, for the actual task, subject performed 90 trials, in six runs of fifteen trials each, separated by a self-paced rest interval. Each run was approximately 6.8 minutes, depending on subjects’ speed (range 4.7–10.7 minutes). Functional data were acquired during these 90 trials.

**Behavior analysis**. For each participant, we extracted the mean reaction time of each conditions (jump to C, jump to E, and no jump). We then performed two-tailed paired t-tests of the mean of jumps against the mean of no jump, and of jump to C against jump to E. We applied the same analysis for accuracy or movement error.

**Image acquisition**. Data were acquired with a 3 T Siemens Allegra (Malvern, PA) head-only MRI scanner, with a circularly polarized head volume coil. High-resolution (1 mm^3^ voxels) T1-weighted structural images were acquired with an MP-RAGE pulse sequence at the beginning of the scanning session. Functional data were acquired using an echo-planar imaging pulse sequence (3 × 3 × 3 mm voxels, 34 contiguous slices, interleaved acquisition, TR of 2000 ms, TE of 30 ms, flip angle 90 º, field of view 192 mm, aligned with the Anterior Commissure – Posterior Commissure plane). The first five volumes of each run were ignored.

**Data analysis**. Data were analyzed using AFNI software (Cox, 1996). The T1-weighted anatomical images were aligned to the functional data. Functional data was corrected for interleaved acquisition using Fourier interpolation. Head motion parameters were estimated and corrected allowing six-parameter rigid body transformations, referenced to the initial image of the first functional run. Data was spatially smoothed with a 6 mm FHWM Gaussian kernel. Each voxels’ signal was converted to percent change.

**General linear model analysis**. For each participant, we created a design matrix modeling events of interest and nuisance variables. At the time of an event of interest we defined an impulse and convolved it with a hemodynamic response. The following regressors were included in the model: (a) an indicator variable marking the occurrence of all pause events, (b) an indicator variable marking the occurrence of jump types E and C, (c) an indicator variable marking the occurrence of type C jumps, (d) a parametric regressor indicating the change in distance to subgoal induced by each C jumps, mean-centered, (e and f) indicator variables marking subgoal and goal attainment, and (g) an indicator variable marking all periods of task performance, from the initial presentation of the icons to the end of the trial. Also included were head motion parameters, first to third order polynomial regressors to regress out scanner drift effects, and global signal, estimated as the mean for each volume. Parameter estimates from the general linear model were normalized to Talairach space (Talairach & Tournoux, 1988), using SPM5 (www.fil.ion.ucl.ac.uk/spm/).

**Group analysis**. For each regressor and for each voxel we tested the sample of 30 subject-specific coefficients against zero in a two-tailed t-test. We defined a threshold of *p* = .01 and applied correction for multiple comparison based on cluster size, using Monte Carlo simulations as implemented in AFNIs AlphaSim. We report results at a corrected *p* < .01.

**Region-of-interest analysis**. We defined nucleus accumbens (NAcc) based on anatomical boundaries on a high-resolution T1-weighted image for each participant. We also defined a region of interest on amygdala using the Talairach atlas in AFNI. Mean coefficients were extracted from these regions for each participant. Reported coefficients for all regions of interest are from general linear model analyses without subtraction of global signal. The sample of 30 subject-specific coefficients from these regions were tested against zero in a two-tailed t-test, with a threshold of *p* < .05.

### Results

**Behavior**. Each trial took 17.16 steps on average across participants (*SEM* = .60 steps). Mean reaction time (RT) for each joystick movement was 1,090 ms (*SEM* = 60 ms). On average, at 3.96 steps (*SEM* =.11 steps), the program interrupted the execution of the task by introducing a pause of 900 ms (which we term pause event, and encompasses the jump and no-jump conditions). In two-thirds of the trials, the envelope jumped to a new location at the onset of the pause (jump condition), and in the remaining third it remained in the same place (no-jump condition). After the pause event was completed, participants took on average 610 ms to produce a new joystick movement (*SEM* = 70 ms; note that there was the enforced delay of 900 ms after for all conditions, which we are not including in our measurement). Participants were significantly slower to react to a jump (*M* = 690 ms, *SEM* = 70 ms) versus a no-jump condition (*M* = 460 ms, *SEM* = 60 ms) as revealed by a two-tailed paired t-test, *t*(29) = 7.96, *p* < .01. However, there was no significant difference between jumps to location C (*M* = 700 ms, *SEM* = 80 ms) *vs*. location E (*M* = 660 ms, *SEM* = 80 ms; *t*(29) = 1.53, *p* =. 14, two-tailed paired t-test).

Mean movement error, measured as the angle between the optimal and the actual joystick movements, across all trial types and participants, was 35.13 º (*SEM* = 1.18 º). Participants were less accurate for movements succeeding a pause event (*M* = 46.37 º*, SEM* = 12.17 º). Movement error for the jump condition (C and E) was higher (*M* = 52.33 º, *SEM* = 28.93 º) than for the no-jump condition (*M* = 33.83 º, *SEM* = 14.55 º; *t*(29) = 6.94, *p* < .01, two-tailed paired t-test). Moreover, participants were less accurate for jump C (*M* = 58.39 º, *SEM* = 39.74 º) than to jump E (*M* = 46.27 º, *SEM* = 22.16 º; *t*(29) = 4.25, *p* < .01, two-tailed paired t-test).

**Whole-brain analysis**. We regressed BOLD activity onto two regressors of interest, a categorical regressor indicating a positive subgoal-related PE (elicited by jumps to location C, see Figure 2) and a parametric regressor for the magnitude of subgoal-related PE (measured as mean-centered decrease in truck-subgoal distance). In the same model, we included three task-specific control regressors (common effect of jump: C + E, mean-centered displacement distance, and common effect of pause event: C + E + no-jump), along with standard control regressors (see Materials and Methods). Complementing the results of Ribas-Fernandes (2011), we observed an increase in BOLD activity in medial prefrontal cortex (mPFC) and right anterior insula to positive subgoal-related prediction errors (cluster-corrected, *p* < .05, Table 1 and Figure 3). Results for control regressors are in Tables 2–5.

**Table 1.** Jumps to C. Experiment I. Whole-brain. Primary threshold p < .01, cluster corrected to p < .05, d.f. 29. Labels provided by Talairach Daemon. Coordinates in Talairach space and DICOM order. G. – gyrus, R. – right, L. – left.

| Area | Size<br>(number of<br>voxels) | Peak voxel |  |  |
| --- | --- | --- | --- | --- |
|  |  | Parameter<br>estimate | t-statistic | Coordinates<br>(X, Y, Z) |
| R. Lingual G. | 462 | -.36 | -5.74 | +0, +71, +1 |
| L. Postcentral G. | 360 | -.25 | -3.41 | +33, +29, +67 |
| R. Superior Frontal G. | 127 | .30 | 3.26 | -33, -46, +31 |
| L. Medial Frontal G.<br>(see Figure 3) | 119 | .14 | 4.62 | +0, -7, +43 |
| L. Lentiform Nucleus | 91 | -.11 | -4.41 | +24, -1, -2 |
| R. Insula | 90 | .12 | 3.03 | -45, -13, +1 |
| L. Medial Frontal G. | 65 | -.11 | -4.26 | +3, +20, +49 |
| R. Lentiform Nucleus | 60 | .09 | -3.80 | -24, -7, +4 |
| L. Middle Frontal G. | 57 | .17 | 3.37 | +36, -28, +43 |
| R. Superior Temporal G. | 53 | -.21 | -2.83 | -51, -19, -20 |
| L. Superior Temporal G. | 53 | -.11 | -4.20 | +54, +20, -2 |

**Table 2.** Decrease in distance to subgoal. Experiment I. Whole-brain. Primary threshold p < .01, cluster corrected to p < .05, d.f. 29. Labels provided by Talairach Daemon. Coordinates in Talairach space and DICOM order. G. – gyrus, R. – right, L. – left.

| Area | Size<br>(number of<br>voxels) | Peak voxel |  |  |
| --- | --- | --- | --- | --- |
|  |  | Parameter<br>estimate | t-statistic | Coordinates<br>(X, Y, Z) |
| L. Precuneus | 1105 | .01 | 5.59 | +0, +77, +49 |
| L. Medial Frontal G. | 118 | -.00 | -3.26 | +0, -61, +19 |
| L. Middle Temporal G. | 113 | -.00 | -4.39 | +66, +5, -11 |
| L. Angular G. | 76 | -.01 | -3.36 | +45, +74, +34 |
| L. Medial Frontal G. | 72 | -.00 | -2.89 | +0, -43, +16 |
| L. Middle Temporal G. | 68 | -.01 | -3.11 | +69, +44, +1 |
| R. Superior Temporal G. | 58 | -.00 | -3.86 | -42, -1, -11 |

**Table 3.** Pause event. Experiment I. Whole-brain. Primary threshold p < .01, cluster corrected to p < .05, d.f. 29. Labels provided by Talairach Daemon. Coordinates in Talairach space and DICOM order. G. – gyrus, R. – right, L. – left.

| Area | Size<br>(number of<br>voxels) | Peak voxel |  |  |
| --- | --- | --- | --- | --- |
|  |  | Parameter<br>estimate | t-statistic | Coordinates<br>(X, Y, Z) |
| R. Superior Temporal G. | 1346 | .37 | 4.91 | -69, +41, +10 |
| L. Precuneus | 1151 | -.37 | -4.04 | +12, +80, +46 |
| L. Precuneus | 1017 | .34 | 5.57 | 0, +59, +43 |
| R. Middle Frontal G. | 882 | -.32 | -3.39 | -36, -43, +34 |
| L. Middle Frontal G. | 835 | -.22 | -4.50 | 24, -19, +55 |
| L. Superior Temporal G. | 798 | .35 | 6.35 | +66, +44, +13 |
| L. Middle Frontal G. | 786 | -.34 | -6.24 | +45, -28, +37 |
| R. Cuneus | 772 | .44 | 2.90 | -3, +95, +13 |
| R. Inferior Parietal Lobule | 703 | -.43 | -4.91 | -51, +44, +52 |
| L. Inferior Frontal G. | 463 | -.34 | -4.58 | +54, -16, -2 |
| L. Postcentral G. | 366 | .29 | 4.47 | +39, +38, +64 |
| R. Medial Frontal G. | 352 | .28 | 3.02 | -3, -61, +4 |
| L. Lentiform Nucleus | 137 | .22 | 6.43 | +21, -4, -2 |
| R. Lentiform Nucleus | 128 | .18 | 4.95 | -21, -4, -2 |
| R. Middle Occipital G. | 124 | -.33 | -3.21 | 33, +95, +4 |
| L. Inferior Temporal G. | 118 | -.21 | -3.27 | +66, +53, -11 |
| R. Middle Frontal G. | 71 | -.12 | -3.93 | -24, +5, +58 |
| R. Caudate Nucleus | 63 | -.16 | -6.06 | -12, +8, +19 |
| L. Caudate Nucleus | 63 | -.16 | -4.29 | +12, +8, +19 |

**Table 4.** Jumps to E and C. Experiment I. Whole-brain. Primary threshold p < .01, cluster corrected to p < .05, d.f. 29. Labels provided by Talairach Daemon. Coordinates in Talairach space and DICOM order. G. – gyrus, R. – right, L. – left.

| Area | Size<br>(number of<br>voxels) | Peak voxel |  |  |
| --- | --- | --- | --- | --- |
|  |  | Parameter<br>estimate | t-statistic | Coordinates<br>(X, Y, Z) |
| R. Precuneus | 839 | .42 | 3.30 | -3, +68, +52 |
| R. Lingual G. | 556 | .25 | 6.09 | 0, +74, +1 |
| R. Insula | 188 | -.11 | -5.42 | -39, +32, +19 |
| L. Fusiform G. | 69 | .13 | 3.47 | +24, +56, -8 |
| L. Superior Frontal G. | 64 | .18 | 3.51 | +30, +8, +64 |
| R. Middle Frontal G. | 58 | -.10 | -3.90 | -39, -31, +16 |

**Table 5.** Displacement distance. Experiment I. Whole-brain. Primary threshold p < .01, cluster corrected to p < .05, d.f. 29. Labels provided by Talairach Daemon. Coordinates in Talairach space and DICOM order. G. – gyrus, R. – right, L. – left.

| Area | Size<br>(number of<br>voxels) | Peak voxel |  |  |
| --- | --- | --- | --- | --- |
|  |  | Parameter<br>estimate | t-statistic | Coordinates<br>(X, Y, Z) |
| L. Precuneus | 228 | .01 | 3.41 | +0, +74, +52 |
| L. Lingual G. | 143 | .00 | 4.38 | +3, +74, +4 |
| R. Traverse Temporal G. | 71 | -.00 | -4.13 | -42, +29, +13 |
| R. Postcentral G. | 59 | -.00 | -3.31 | -60, +20, +40 |

**Figure 3.**
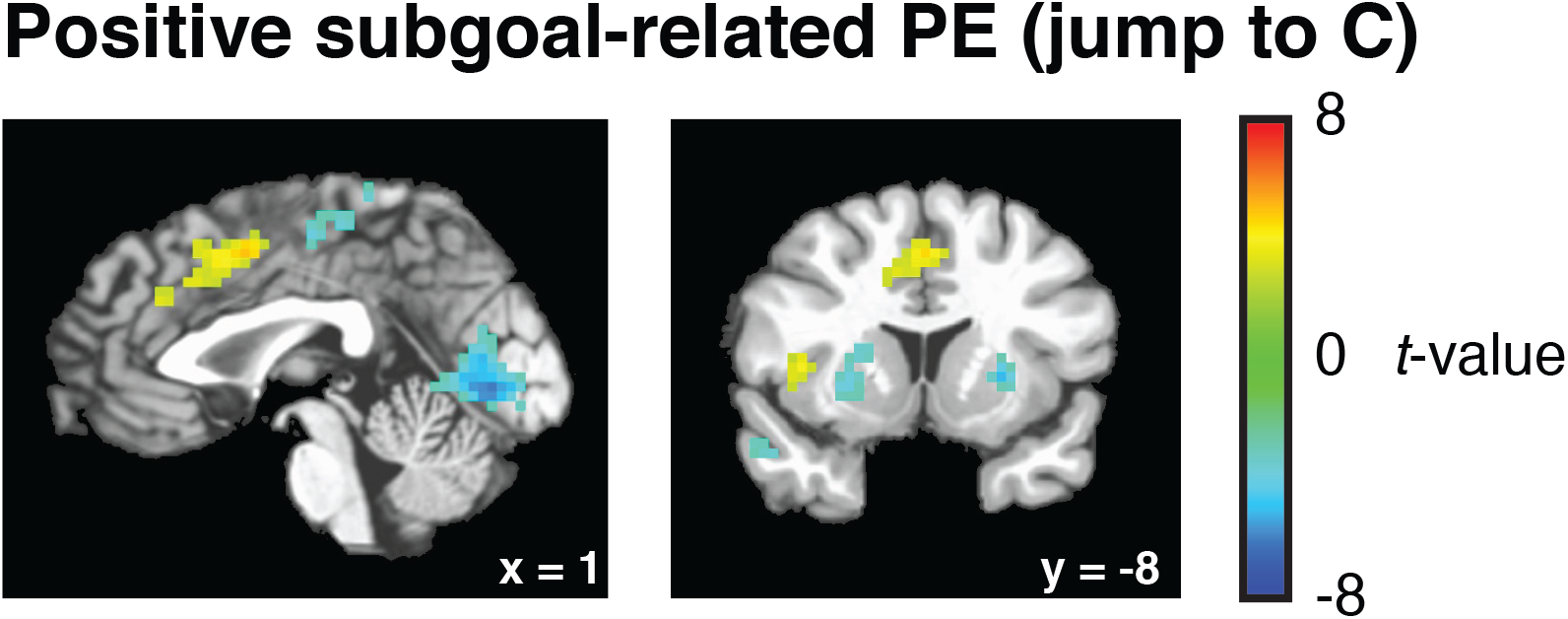
Whole-brain results of Experiment I, jump to C. Jumps that featured a decrease in distance to the envelope, without changing the overall distance to the house, were associated with an increase in BOLD activity in medial PFC and anterior insula. This effect is independent from spatial reorientation, as suggested by the absence of activity in these areas to jumps to E, a condition with the highest angle of displacement and no changes in distance to the envelope.

**Region-of-interest analysis**. To investigate whether areas known to process goal-related PEs were responsive to subgoal-related PEs in our experiment, we anatomically delineated two regions of interest, NAcc, and amygdala.

No significant change in BOLD response was observed to jumps to point C in anatomically defined bilateral NAcc (mean of regression coefficient, *M* = −1.83 × 10^−3^, *p* = .94). Qualitatively similar results were obtained when the same analysis was performed on left and right NAcc separately. Similar null results, bilaterally and unilaterally, were observed in the amygdalar region (Jump to C, *M* = -.04, *p* = .06; parametric decrease in subgoal distance, *M* = -3.58 × 10^−4^, *p* = .30).

## Experiment II: An Examination of Goal and Subgoal-Related PEs

To recap, our paradigm elicits different types of prediction errors by having the subgoal unexpectedly jump to different points in space (as illustrated in Figure 2). In Experiment II, similar to Experiment I, two-thirds of the trials featured a jump to a new location, whereas in the remaining third the location of the envelope did not change. However, in contrast with Experiment I, all of the jumps were to the locations that should putatively elicit both goal-related PEs and subgoal-related PEs. We manipulated the displacement of the jumps so that the magnitude of different types of prediction errors was uncorrelated across the experiment. We opted for a parametric design rather than a categorical design, because the categorical manipulation of types and valence of prediction errors would have resulted in a high number of conditions: four experimental conditions, A, B, C, D and two control conditions, E and no-jump, which would yield a lower number of trials per condition.

As an additional modification of Experiment I, we included goal-related PEs driven by probabilistic monetary rewards and punishments, at the end of each trial (see Figure 4, and Materials and Methods). This was introduced to compare the correlates of distance-driven goal-related PEs in our experiments with those of more standard monetary goal-related PEs.

**Figure 4.**
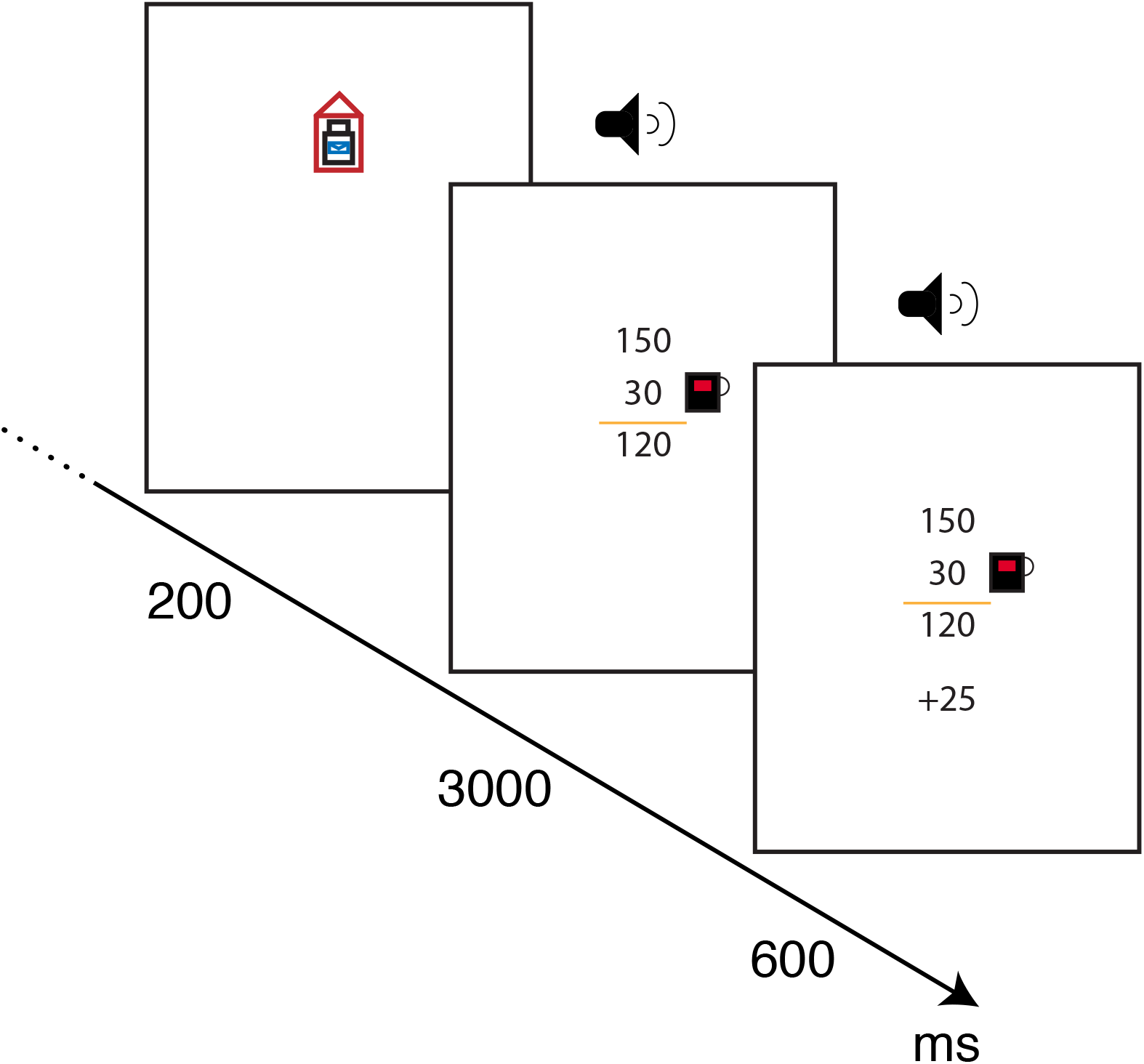
Eliciting monetary prediction errors. In the second experiment, at the end of each trial, participants would receive information about their performance. A delivery yielded 150 points, and any additional step from the shortest distance possible would be deducted from this rate. In the example above, 30 points were deducted for extra steps. In addition, a probabilistic outcome was introduced (+25). Unrelated to their performance, participants would receive a bonus of 25 points, 0 points or −25 points, with equal probability. Points accrued would be exchanged for dollars at the end of the experiment.

Furthermore, we deducted errors in performance from the monetary bonus at the end of each trial (see Figure 4). This was done to ensure that participants were highly motivated to reach the overall goal in the lowest number of steps.

### Materials and Methods

**Participants**. Forty-eight participants were recruited from the Princeton University community, and 8 participants were excluded, 7 for having incidents of head movements larger than 2.5 mm and 1 for failure to complete the task inside the scanner (ages 18–27 years, *M* = 20, 15 male, 38 were right-handed and 2 were left-handed, joystick was always held in the right hand). All participants received monetary compensation at a departmental standard rate. In addition, participants received two types of monetary bonuses, one based on performance, and a probabilistic “tip”, as described below.

**Materials, task and procedure**. Similar to Experiment I, the task consisted of three parts: a short behavioral practice outside the scanner, for 12 trials, using a joystick held in the right hand (Logitech International, Romanel-sur-Morges, Switzerland), a 12 trial practice inside the scanner, using an MR compatible joystick (MagConcept, Redwood City, CA) during structural scan acquisition, and a third phase of 132 trials (6 runs of 22 trials) for approximately sixty minutes (run duration, *M* = 11.7 minutes, *SEM* = .3 minutes), where functional data were collected. Each run started and ended with a central fixation cross, displayed for 10,000 ms. At the end of each run participants would be given a self-paced break.

On each trial, the house occupied the same vertex as in Experiment I (0,-200). However, the initial position of the truck and the envelope were different and determined as follows. The initial position of the truck (-90,320) was set so that it would be 150 pixels, or 3 optimal steps, from a virtual line beyond which a jump would be triggered. This line was parallel to the house-envelope line and would go through the point (0,200), a point where the envelope is at the same distance to the house and to the truck. This location is convenient because it allows for equal variance in both positive and negative prediction errors.

Similar to Experiment I, when a jump was triggered, a brief tone was played, the truck and the envelope would flash yellow, and joystick movements were ignored for 900 ms. This pause happened on average after 5.6 steps (*SEM* = .1 steps). In one-third of the trials (44) the envelope would stay in the same location. In the remaining two-thirds (88) it would jump to a new location (see the next paragraph for details on the jump locations). Instructions to participants were the same as in Experiment I.

In this experiment, each jump generates a goal-related PE, a subgoal-related PE, and requires spatial and motor coordination that is proportional to the nuisance variable displacement distance. We applied a Monte Carlo approach to obtain different sets of 88 jump locations for each participant in order to decorrelate the variables goal-related PE magnitude, subgoal-related PE magnitude, and displacement distance. We sampled the space of (x,y) coordinates with the restrictions that there should be an equal number of positive and negative goal-related PEs and subgoal-related PEs, an equal number of ipsilateral and contralateral jumps, and that negative goal-related PEs should have a similar range to that of positive goal-related PEs. After sampling within these regions, for each participant we selected sets of 88 jumps that (1) had a mean subgoal-related PE close to zero (less than half a standard deviation away from zero), (2) had a mean goal-related PE close to zero (less than a third of a standard deviation away from zero), (3) had a low sum of correlation between variables, and (4) had a high variance. As mentioned before, due to variability in performance (see accuracy in Experiment I), we predicted there would be variability in the locations that triggered a jump. In the observed behavioral data, the correlation between subgoal-related PE and goal-related PE was .31, correlation between subgoal-related PE and displacement distance 0, correlation between goal-related PE and displacement distance -.37, and the average goal-related PE and subgoal-related PE were close to zero (*M* = .10 and .00 steps respectively).

As in Experiment I, after the jump, participants headed towards the new location of the subgoal. When the truck passed within 30 pixels of the envelope, the envelope moved to the truck and remained there for the subsequent moves. When the truck with the envelope passed within 30 pixels of the house, the truck with the envelope appeared within the house. This image was displayed for 200 ms.

Participants were paid at a rate of 150 points per delivery, a task currency that would be converted to US dollars. Though they were not told what the conversion rate was, they were told that “if they worked hard” a maximum of $12 could be attained at the end of Experiment In addition to the departmental rate. As mentioned before, we deducted errors in performance from the monetary bonus for each delivery at the rate of .1 points *per* pixel travelled, up to a maximum of 100 points. We presented this information for 3,000 ms, in a screen after the truck entered the house (see Figure 4). This was accompanied by the sound of cash register.

After 3,000 ms, a probabilistic monetary reward appeared at the bottom of the screen (Figure 4). This was introduced to compare distance-driven goal-related PEs with monetary goal-related PEs. Participants could get 25, -25 or 0 points to points with equal probability. They were told that this was not contingent on their performance but that it was worthwhile to pay attention to this additional payment, given that final payment was a sum of rewards accrued during all task phases. To ensure attentional capture, we introduced a sound at the moment of this information (coin sound for 25, different from the one for the at rate, a sad trumpet sound for -25, and a brief tone for 0, all sounds had the same 100 ms duration). This probabilistic monetary reward was displayed for 600 ms and was followed by a fixation cross that remained on screen for 700 ms.

**Image acquisition**. Data were acquired with a 3T Siemens Skyra (Malvern, PA) MRI scanner using a sixteen-channel head coil. High resolution (1 mm3 voxels) T1-weighted structural images were acquired with an MP-RAGE pulse sequence at the beginning of the scanning session.

Functional data were acquired using a high-resolution echo-planar imaging pulse sequence (3 × 3 × 3 mm voxels, 35 contiguous slices, 3 mm thick, interleaved acquisition, TR of 2,000 ms, TE of 30 ms, flip angle 90 º, field of view 192 mm, aligned with the Anterior Commissure - Posterior Commissure plane). The first five volumes of each run were ignored.

**Data analysis**. Data were analyzed using AFNI software (Cox, 1996). The T1-weighted anatomical images were aligned to the functional data. Functional data was corrected for interleaved acquisition using Fourier interpolation. Head motion parameters were estimated and corrected allowing six-parameter rigid body transformations, referenced to the initial image of the first functional run. A whole-brain mask for each participant was created using the union of a mask for the first and last functional images. Spikes in the data were removed and replaced with an interpolated data point. Data was spatially smoothed with a 6 mm FHWM Gaussian kernel. Each voxels signal was converted to percent change by normalizing it based on intensity.

**General linear model analysis**. For each participant, we created a design matrix modeling experimental events and including events of no interest. At the time of an experimental event we defined an impulse and convolved it with a hemodynamic response. The following regressors were included in the model: (a) an indicator variable marking the occurrence of all auditory tone / envelope events, (b) an indicator variable marking the occurrence of all jump events, (c) a parametric regressor indicating the change in distance to subgoal induced by each jump, mean-centered, (d) a parametric regressor indicating the change in distance to goal induced by each jump, mean-centered, (e and f) indicator variables marking subgoal and goal attainment, (g) a variable marking all periods of task performance, from the initial presentation of the icons to the end of the trial, (h) an indicator variable for delivery of monetary reward (encompassing the positive, 25, negative, -25, and neutral, 0, events), (i) an indicator variable for the positive reward, 25, and (j) an indicator variable for the negative reward, -25. Also included were head motion parameters, and first to third order polynomial regressors to regress out scanner drift effects. A global signal regressor was also included. In additional analyses, instead of indicator variables encompassing signed positive and negative events, we separated regressors for positive and negative events, or included them in an unsigned way, with one regressor for the jump PEs and one regressor for the monetary PEs. All parametric regressors were mean-centered. The estimates from the general linear model were normalized to Talairach space (Talairach & Tournoux, 1988).

**Group analysis**. For each regressor and for each voxel we tested the sample of 40 subject-specific coefficients against zero in a two-tailed t-test. We defined a threshold of *p* = .01 and applied correction for multiple comparison based on cluster size, using Monte Carlo simulations as implemented in AFNIs AlphaSim. We report results at a corrected *p* < .01.

**Region-of-interest analysis**. We defined ventral striatum (including the olfactory tubercle) based on individual anatomical boundaries. We also defined a region of interest on the amygdala based on normalized Talairach atlas. Mean coefficients were extracted from this region for each participant. Reported coefficients for all regions of interest are from general linear model analyses without subtraction of global signal. The sample of 40 subject specific coefficients were tested against zero in a two-tailed t-test, with a threshold of *p* < .05.

### Results

**Behavior**. A trial lasted on average 19.81 steps (*SEM* = .40 steps). Average RT was 1,160 ms (*SEM* = 30 ms). The pause happened on average at 5.57 steps (*SEM* =. 08 steps). Average RT for the first movement after pause events was 1,460 ms (*SEM* = 30 ms). A linear regression of RTs of pause events revealed a significant increase in RTs to jumps (*mean regression coefficient* = .07, *SEM* = .02; *t*(39) = 3.28, *p* < .005, two-tailed t-test). The same regression also revealed that RTs were significantly slower as displacement distance increased (*mean regression coefficient* = .04, *SEM* = .01; *t*(39) = 4.22, *p* < .001). No significant effect of subgoal-related PE or goal-related PE was observed (*p* = .15 and *p* = .78). Similar results were obtained with mean-centered unsigned regressors for both subgoal-related PEs and goal-related PEs.

Mean accuracy, across all steps, was 71.68 º (*SEM* = .21 º), and, for the step immediately after the pause event, the accuracy was 35.69 º (*SEM* = 1.1 º). A linear regression on the accuracy scores on the step succeeding the pause event revealed a significant increase in deviations from the optimal path in the jump condition (*mean regression coefficient* = .08, *SEM* =. 02; *t*(39) = 4.43, *p* < .05). We also observed that the extent of deviation from optimal path increased with displacement distance (*mean regression coefficient* =. 03, *SEM* = .01; *t*(39) = 2.11, *p* < .05). Similar with what we observed with RT data, no significant effect of subgoal-related PEs or goal-related PEs was observed (*p* = .25 and *p* = .11).

**Whole-brain analysis**. We observed an increase in BOLD response in left mPFC to distance-driven unsigned goal-related prediction errors (*M* = 1.0 × 10^−3^, *p* < .05, cluster corrected; Figure 5 and Table 6). Surprisingly, in contrast with the previous experiment, and our past study, no response was observed to unsigned subgoal-related prediction errors, even at a liberal threshold (Table 7). Regression models with signed regressors yielded results consistent with the models with unsigned responses. Results for control regressors are in Tables 8 to 10.

**Figure 5.**
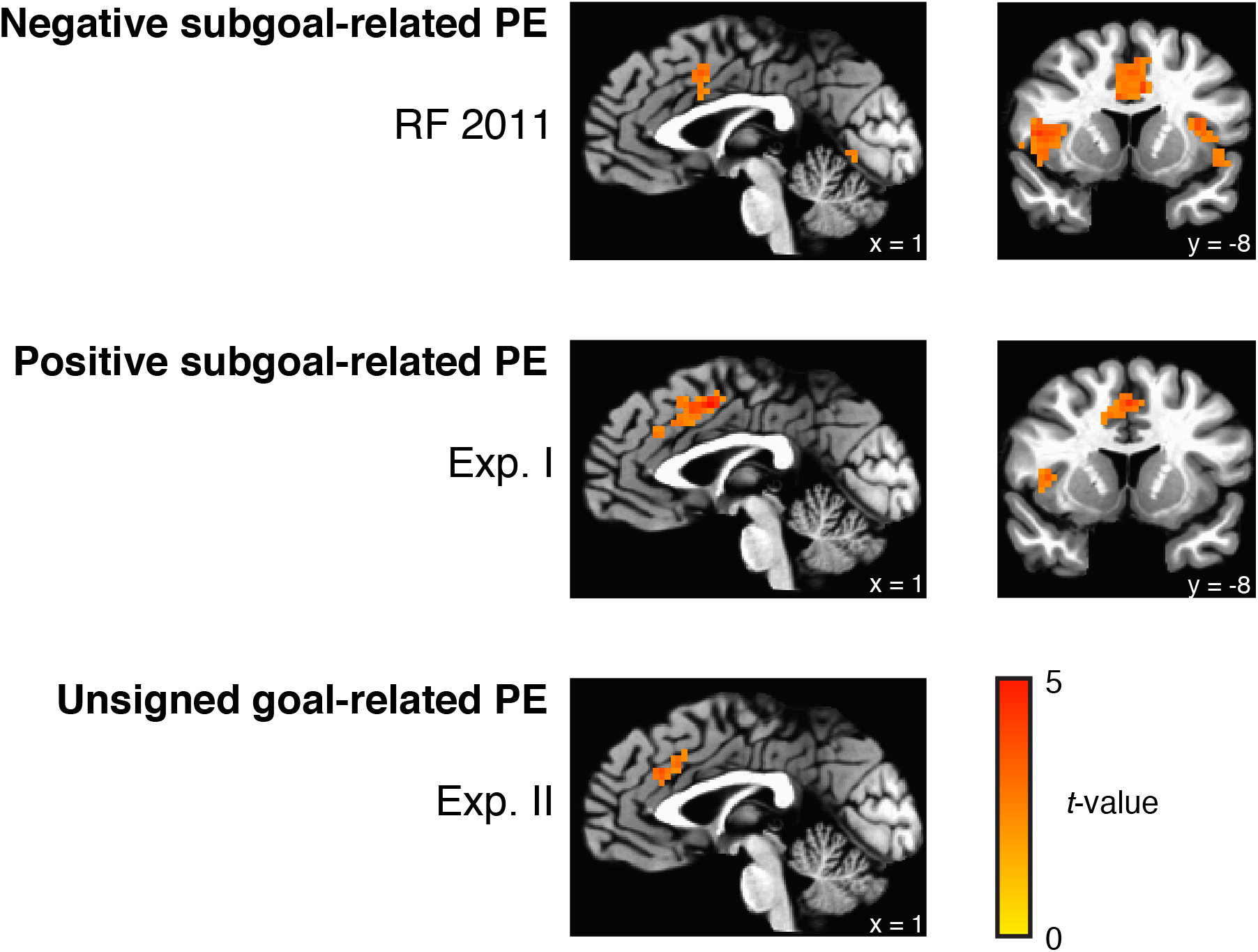
Comparison of unsigned responses across Experiments I, II and Ribas-Fernandes, et al. (2011). For comparison, only the positive clusters in Experiment I are shown—see Figure 1 for an image with positive and negative clusters.

**Table 6.** Unsigned distance-driven goal-related PE. Experiment II. Whole-brain. Primary threshold p < .01, cluster corrected to p < .05, d.f. 29. Labels provided by Talairach Daemon. Coordinates in Talairach space and DICOM order. G. – gyrus, R. – right, L. – left.

| Area | Size<br>(number of<br>voxels) | Peak voxel |  |  |
| --- | --- | --- | --- | --- |
|  |  | Parameter<br>estimate | t-statistic | Coordinates<br>(X, Y, Z) |
| L. Cingulate G. | 45<br>($p = .07$) | .00 | 4.20 | +1, -28, +29 |

**Table 7.**
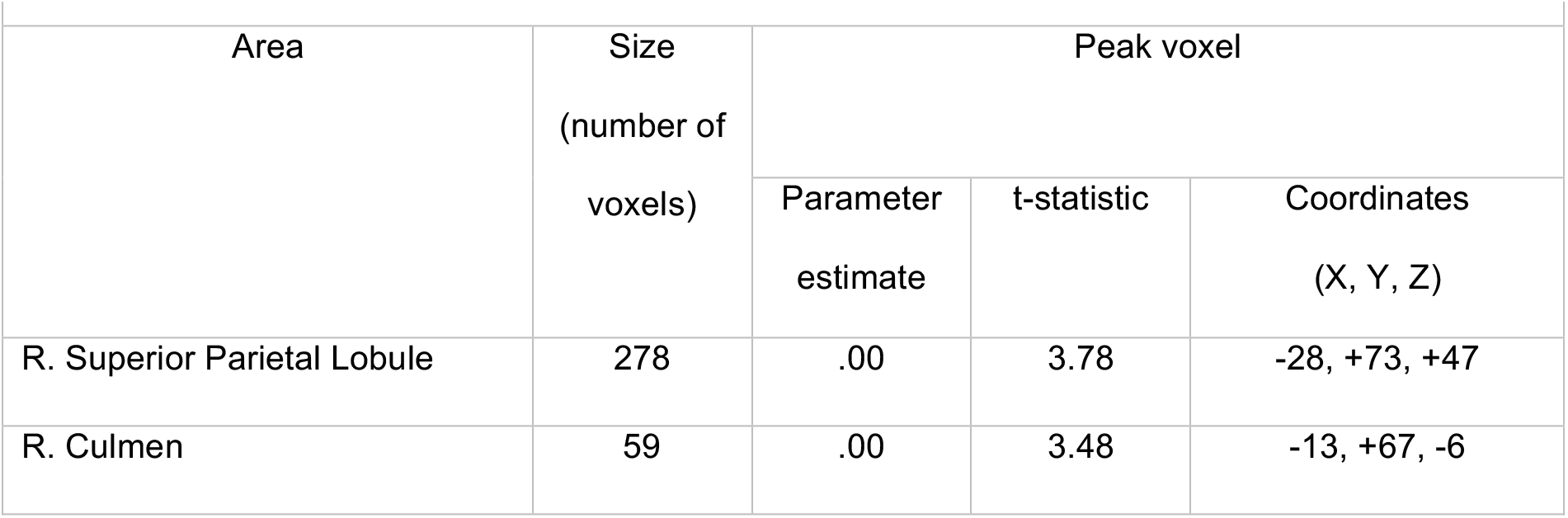
Unsigned subgoal-related PE. Experiment II. Whole-brain. Primary threshold p < .01, cluster corrected to p < .05, d.f. 29. Labels provided by Talairach Daemon. Coordinates in Talairach space and DICOM order. G. – gyrus, R. – right, L. – left.

**Table 8.** Pause event. Experiment II. Whole-brain. Primary threshold p < .01, cluster corrected to p < .05, d.f. 29. Labels provided by Talairach Daemon. Coordinates in Talairach space and DICOM order. G. – gyrus, R. – right, L. – left.

| Area | Size<br>(number of<br>voxels) | Peak voxel |  |  |
| --- | --- | --- | --- | --- |
|  |  | Parameter<br>estimate | t-statistic | Coordinates<br>(X, Y, Z) |
| R. Superior Frontal G. (cluster<br>extends to anterior medial<br>surface) | 9555 | .40 | 6.00 | -1, -37, +56 |
| L. Inferior Parietal Lobule | 8646 | -.46 | -5.55 | +46, +46, +53 |
| L. Superior Frontal G. | 1642 | -.41 | -0.60 | +37, -40, +32 |
| R. Middle Frontal G. | 686 | -.340 | -3.84 | -34, -43, +35 |
| R. Inferior Frontal G. | 684 | -.28 | -5.69 | -49, -16, 0 |
| L. Cuneus | 468 | .43 | 5.37 | +1, +9, +5 |
| L. Inferior Temporal G. | 63 | -.13 | -4.19 | +55, +58, -3 |

**Table 9.** Displacement distance. Experiment II. Whole-brain. Primary threshold p < .01, cluster corrected to p < .05, d.f. 29. Labels provided by Talairach Daemon. Coordinates in Talairach space and DICOM order. G. – gyrus, R. – right, L. – left.

| Area | Size<br>(number of<br>voxels) | Peak voxel |  |  |
| --- | --- | --- | --- | --- |
|  |  | Parameter<br>estimate | t-statistic | Coordinates<br>(X, Y, Z) |
| R. Superior Frontal G. | 106 | .00 | 2.90 | -28, -4, +65 |
| L. Inferior Parietal Lobule | 96 | .00 | 4.26 | +52, +43, +53 |
| L. Superior Temporal G. | 81 | .00 | 2.86 | -49, -10, +2 |
| R. Precuneus | 71 | .00 | 3.11 | -1, +55, +47 |
| L. Postcentral G. | 63 | -.00 | -3.33 | +28, +31, +65 |
| L. Superior Frontal G. | 59 | .00 | 3.55 | +37, -52, +26 |
| R. Supramarginal G. | 50 | -.00 | 3.49 | -61, +46, +29 |
| R. Superior Temporal G. | 49 | -.00 | -3.42 | -52, +28, +14 |

**Table 10.** Jumps A, B, C, and D. Experiment II. Whole-brain. Primary threshold p < .01, cluster corrected to p < .05, d.f. 29. Labels provided by Talairach Daemon. Coordinates in Talairach space and DICOM order. G. – gyrus, R. – right, L. – left.

| Area | Size<br>(number of<br>voxels) | Peak voxel |  |  |
| --- | --- | --- | --- | --- |
|  |  | Parameter<br>estimate | t-statistic | Coordinates<br>(X, Y, Z) |
| R. Precuneus | 4438 | .43 | 5.41 | -10, +73, +53 |
| L. Precentral G. | 925 | -.14 | -3.18 | +58, -4, +5 |
| L. Middle Frontal G. | 758 | .19 | 4.07 | -19, -10, +62 |
| L. Anterior Cingulate | 717 | -.18 | -3.43 | +1, -16, -6 |
| L. Precentral G. | 398 | -.15 | -5.06 | +34, +19, +65 |
| R. Middle Frontal G. | 222 | .12 | 3.76 | -31, -34, +41 |
| L. Inferior Frontal G. | 198 | -.15 | -2.71 | +43, -25, -12 |
| L. Uncus | 194 | -.11 | -3.78 | +16, +7, -21 |
| R. Transverse Temporal G. | 165 | -.10 | -3.59 | -64, +13, +11 |
| L. Middle Frontal G. | 139 | .13 | 4.25 | +40, -31, +38 |
| L. Declive | 121 | -.09 | -2.87 | +13, +73, -15 |
| L. Caudate | 119 | .07 | 5.34 | +16, -10, +8 |
| L. Parahippocampal G. | 98 | .07 | 5.52 | +28, +46, -6 |
| L. Cingulate G. | 89 | -.07 | -3.47 | +1, +25, +41 |
| L. Postcentral G. | 84 | -.13 | -3.38 | +1, +49, +23 |
| R. Middle Temporal G. | 76 | -.12 | -2.99 | -61, +4, -21 |
| R. Inferior Frontal G. | 70 | -.13 | -3.21 | -40, -22, -12 |
| R. Parahippocampal G. | 68 | -.08 | -2.91 | -16, +4, -9 |
| R. Superior Temporal G. | 62 | -.08 | -2.86 | -58, -1, +5 |
| R. Caudate | 58 | .04 | 4.76 | -16, -7, +14 |
| R. Parahippocampal G. | 53 | .05 | 3.63 | -31, +34, -9 |
| L. Precuneus | 48 | -.07 | -3.08 | +1, +70, +23 |

After each delivery, a probabilistic monetary reward was delivered: +25, 0, -25, with equal probability. In contrast with the unsigned distance-driven reward prediction error, we observed no medial prefrontal activity to signed or unsigned monetary prediction errors (+25, -25), compared with (+25, 0, -25), even at liberal thresholds – see Table 11. Positive monetary reward prediction errors (+25) yielded an increase BOLD response in left putamen activity, on the border between ventral and dorsal striatum (see Figure 6 and Table 12 for coordinates), relative to the common responses to all the possible monetary outcomes, +25, 0, -25. This was matched by a contralateral striatal cluster at a more liberal threshold. In addition, we observed increases in bilateral fusiform gyrus and a decrease in response in bilateral superior temporal gyrus. Contrary to our expectations, no changes in medial prefrontal signal were observed to monetary reward prediction errors.

**Figure 6.**
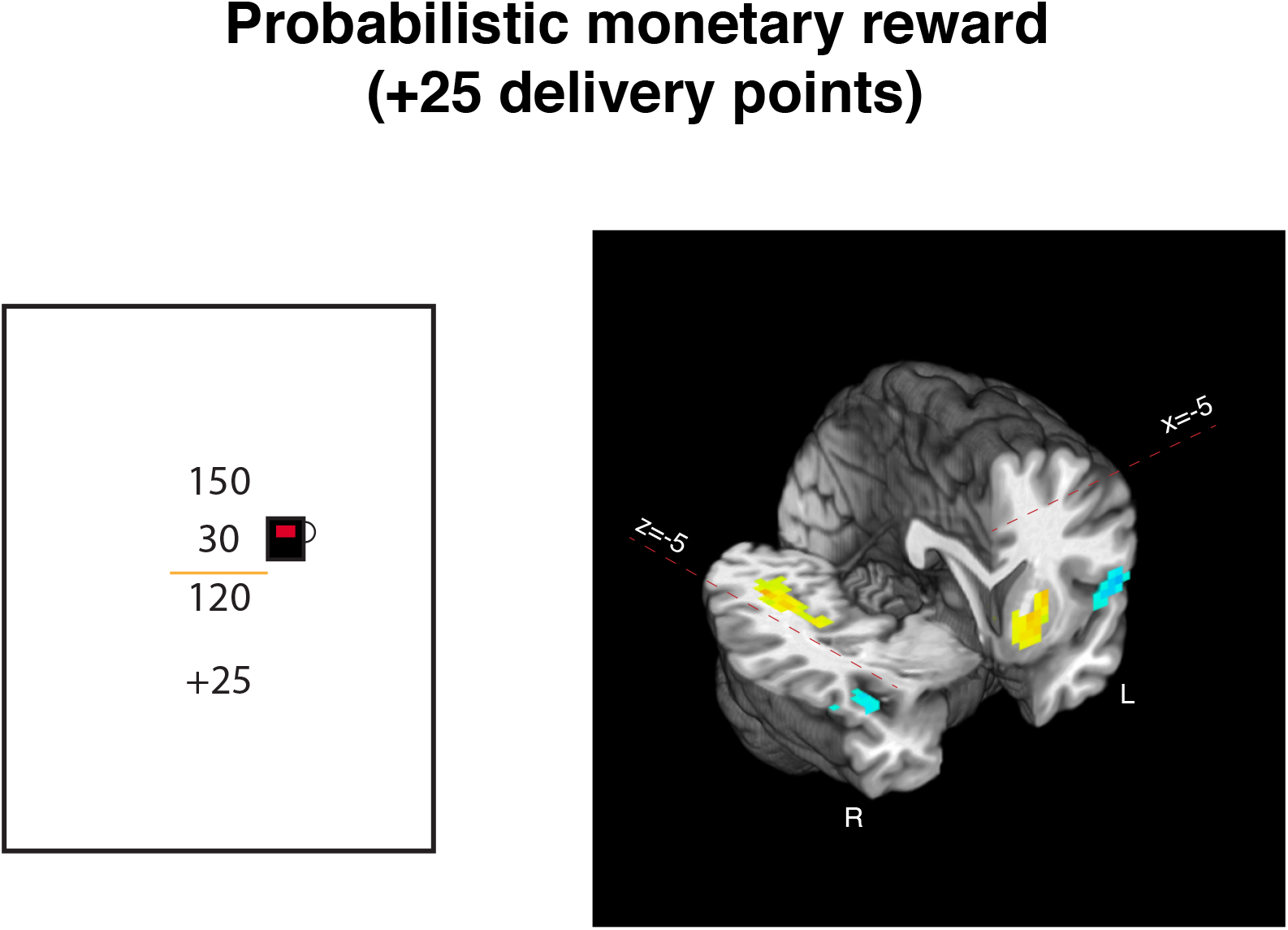
Eliciting positive reward prediction errors with monetary outcomes. In the second experiment, at the end of each trial, in addition to performance based reward (see Figure 4). On a third of trials participants would get +25 delivery points, which would later be converted to dollars. We observed left ventral putamen increases to tip, compared with outcome (+25, 0 or −25). In addition, we observed bilateral decreases in response in superior temporal gyrus, and increases in fusiform gyrus. *p* < .05, cluster corrected (see Table 12 for coordinates).

**Table 11.**
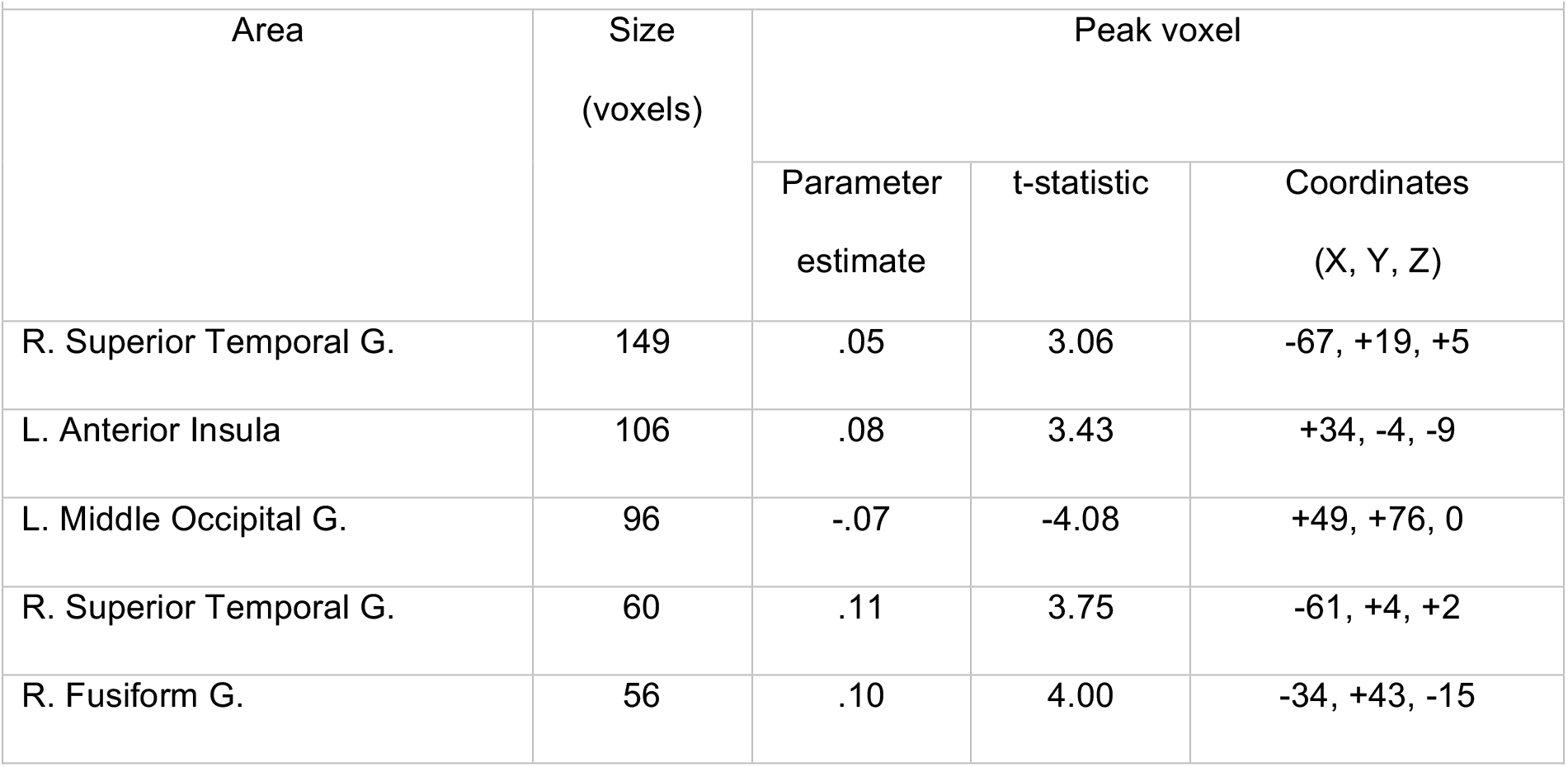
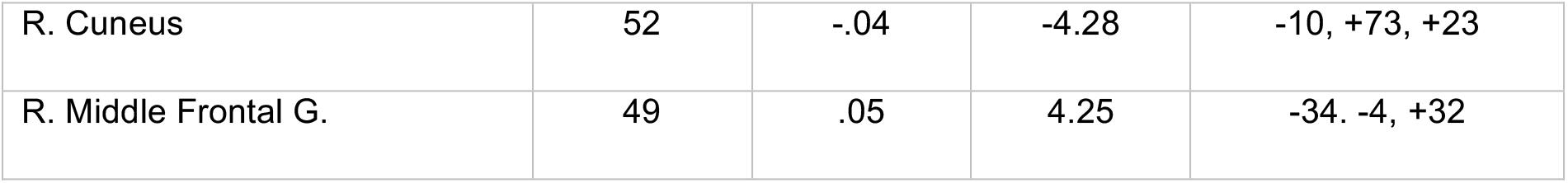
+25 and −25, compared with +25, 0 and −25 points (unsigned monetary probabilistic reward independent of delivery; see Figure 6). Experiment II. Whole-brain. Primary threshold p < .01, cluster corrected to p < .05, d.f. 29. Labels provided by Talairach Daemon. Coordinates in Talairach space and DICOM order. G. – gyrus, R. – right, L. – left.

**Table 12.** +25 points (monetary probabilistic reward independent of delivery; see Figure 6). Experiment II. Whole-brain. Primary threshold p < .01, cluster corrected to p < .05, d.f. 29. Labels provided by Talairach Daemon. Coordinates in Talairach space and DICOM order. G. – gyrus, R. – right, L. – left.

| Area | Size<br>(number of<br>voxels) | Peak voxel |  |  |
| --- | --- | --- | --- | --- |
|  |  | Parameter<br>estimate | t-statistic | Coordinates<br>(X, Y, Z) |
| L. Superior Temporal G. | 367 | -.24 | -4.32 | +64, +25, +14 |
| R. Superior Temporal G. | 352 | -.27 | -5.01 | -64, +13, +8 |
| R. Fusiform G. | 201 | .09 | 2.81 | -40, +49, -21 |
| L. Fusiform G. | 94 | .08 | 5.69 | +28, +49, -12 |
| L. Lentiform Nucleus | 41 | .10 | 3.65 | +16, -1, -6 |

**Region-of-interest analysis**. No significant response was observed in anatomically delineated ventral striatum to subgoal-related PEs, or distance-driven goal-related PEs (*p* > .05). Consistent with whole-brain results, we did observe a significant response in the bilateral putamen, on the border with ventral striatum, to monetary positive reward prediction errors (*p* < .001).

We tested for unsigned reward prediction errors in anatomically defined amygdalar complex. We found no significant response to distance-driven goal-related PEs, subgoal-related PEs, or to monetary goal-related PEs (*p* > .05).

## Discussion

To explore the neural correlates of goal-related PEs and subgoal-related PEs we conducted three experiments manipulating negative subgoal-related PEs (Ribas-Fernandes et al., 2011), positive subgoal-related PEs (Experiment I), and negative/positive subgoal-related PEs, together with goal-related PEs (Experiment II). We found that mPFC responded in an unsigned manner to subgoal-related PEs (Experiment I and RF2011). Unsigned responses in mPFC have been found consistently to reward prediction errors, both in electrophysiological and neuroimaging studies (Bryden, Johnson, Tobia, Kashtelyan, & Roesch, 2011; Hayden, Heilbronner, Pearson, & Platt, 2011; Hyman et al., 2017; Roesch et al., 2012). Multiple theories of mPFC function, which includes dorsal anterior cingulate cortex predict an unsigned BOLD response to prediction errors, namely Expected Value of Control (Shenhav, Botvinick, & Cohen, 2013), Predicted Response-Outcome (Alexander & Brown, 2011), Attention to Learning theory of ACC (Roesch et al., 2012), Reward Value and Prediction Model (Silvetti, Seurinck, & Verguts, 2011).These theories contrast with other theories predicting signed modulation of BOLD response (Holroyd & Coles, 2012). Regardless of a specific theoretical framework, our findings suggest that the role of mPFC extends to decision variables related to subtask performance.

Consistent with previous findings (Ribas-Fernandes et al., 2011), mPFC activity correlated with PEs was found to be specific to changes in distance to goal and subgoal and could not be explained away by geometric changes associated with a jump. We can exclude geometric factors because our PE findings are relative to a control condition that manipulated displacement distance of the envelope, without changing distance to the subgoal or goal (jump E). To further rule out a geometric account of mPFC activity, we included a regressor of displacement distance, which did not reveal any response in mPFC (Table 5).

As mentioned before, Experiment II simultaneously manipulated sub-goal related PEs, goal-related PEs, and monetary PEs. However, in striking contrast to Experiment I and Ribas-Fernandes et al. (2011), no subgoal-related PEs were observed in Experiment II. We expected mPFC in Experiment II to show unsigned subgoal-related PEs, in keeping with the previous experiments, independently of unsigned goal-related PEs. Yet, we only observed the unsigned goal-related PEs.

Importantly, the modulations of BOLD response in our study cannot be explained away by the mere change in allocation of attentional resources resulting from the displacement of the envelope, as we did not observe any parametric manipulation of the mPFC response by displacement distance. In other words, the mPFC’s responses unequivocally reflected the evaluation of the jump event in relation to the goal and subgoal state

It is noteworthy that in Experiment II, subgoal and goal-related PEs were induced simultaneously by displacing the subgoal. This raises the possibility that when information about goals and subgoals is presented simultaneously, humans actively disregard information pertaining to lower level of task hierarchy, i.e., subgoals, in favor of goal-related information. Another non-mutually exclusive possibility is that the capacity to compute two prediction errors from one source of information in this task is limited, thus one of the levels may not be processed properly. A possible candidate for such difficulty might be the complexity to process local and global distances simultaneously in this task. Accordingly, these findings suggest a new principle for mPFC PE signaling in hierarchically structured tasks. Specifically, the data are consistent with a critical role for attention, whereby PE signals are generated based on the specific level of task structure that is currently attended, as determined by the specific contingencies of the task. While mPFC was sensitive to distance-driven PEs (subgoal-related PEs and goal-related PEs), VS was only sensitive to money-driven goal-related PEs (Experiment II). The differential engagement of mPFC and VS is compatible with the hierarchical reinforcement learning theory of mPFC function (Holroyd & Yeung, 2012). The theory contends that mPFC is highly engaged in tasks that require extended sequences of actions and involve effortful behavior. Our spatial navigation task incorporates both features, where each trial was composed a series of 17 to 19 mentally effortful joystick movements (see Behavioral Results for both experiments). Another prominent theory of mPFC function posits that this region is involved in the evaluation of the expected value of control (Shenhav et al., 2013). In our task, it is conceivable that participants had to overcome the impulse to move the joystick in the direction of the current destination, in order to displace the truck in an optimal way. The process of overriding prepotent responses is known to require control (Cohen, Dunbar, & McClelland, 1990; Ridderinkhof, Ullsperger, Crone, & Nieuwenhuis, 2004). Hypothetically, any displacement in the subgoal changes the level of control required to complete a delivery. From Shenhav et al. (2013), it follows that mPFC should be parametrically sensitive to such displacements.

It is interesting to compare our findings with the results of Diuk et al. (2013) where ventral striatum was sensitive to simultaneous prediction errors at two different levels of task hierarchy. In Diuk et al., the information pertinent to task levels were presented explicitly as two different stimuli, whereas in our study such information should be inferred by attending to the change in the relative arrangement of stimuli resulting from jump displacement. Taken together, the results of Diuk et al. (2013) and Experiment II suggest that humans can be sensitive to two sources of information simultaneously provided these sources are presented separately.

In conclusion, our results indicate that mPFC has a role in the processing of the information that are hierarchically structured. More specifically, we show that (1) mPFC signals subgoal-related PEs in an unsigned manner, (2) mPFC signals PEs related to superordinate goals similarly, (3) whether mPFC’s BOLD response reflects subgoal or goal-related PE is dependent on the specific task manipulation and is presumably determined by attentional factors. (4) PE signaling differs between mPFC and VS. Such prediction errors are presumably used to improve behavior at the level of subtasks, which can then be applied to different tasks. Given that ecological tasks are hierarchically structured, mPFC can be instrumental in extending reinforcement learning mechanisms to ecological settings.

